# Photo-Electrochemical Stimulation of Neurons with Organic Donor-Acceptor Heterojunctions

**DOI:** 10.1101/2022.02.17.480608

**Authors:** Achilleas Savva, Adel Hama, Gabriel Herrera-López, Nicola Gasparini, Ludovico Migliaccio, Malak Kawan, Nadia Steiner, Iain McCulloch, Derya Baran, Hubert Fiumelli, Pierre Magistretti, Eric D. Głowacki, Sahika Inal

## Abstract

Recent advancements in light-responsive materials enabled the development of devices to artificially activate tissue with light, and show great potential for use in different types of therapy. Photo-stimulation based on organic semiconductors has recently attracted interest due to their unique set of properties such as biocompatibility, better mechanical match with human tissue, and strong absorption of light in the visible spectrum. Here we show the development of solution processed organic heterojunctions that are able to control the activity of primary neurons *in vitro* with light. The p-type polymer semiconductor PDCBT and the n-type polymer semiconductor ITIC (also known as non-fullerene acceptor) are simply spin coated on glass substrates forming a bilayer p-n junction with high photo-sensitivity in aqueous electrolytes. Photo-electrochemical measurements reveal that high photo-voltage and photo-current is produced, as a result of a charge transfer between the polymers and oxygen in the electrolyte. The biocompatibility of the proposed materials is addressed with live/dead assays on both primary mouse cortical neurons and human cell lines that are cultured on their surface. We have found that light of low intensity (i.e. 40 mW/cm^2^) is absorbed, and converted into a cue that triggers action potential on primary cortical neurons directly cultured on glass/PDCBT/ITIC interfaces as proven by patch clamp measurements. The activation of neurons is most likely due to photochemical reactions at the polymer/electrolyte interface that result in hydrogen peroxide, which might lead to modulation of specific ion channels on neurons membrane. Photo-thermal effects are excluded with controlled patch clamp measurements on neurons cultured on plain glass and on photoresist thin films. The profound advantages of low intensity light stimulation, simplified fabrication, and wireless operation pave the way for the integration of these interfaces in multiplex bioelectronic devices for the development of novel light therapy concepts and powerful neuroscience research tools.

## Introduction

Biomedical engineering concepts that use light to control cellular activity have been widely used in clinical practice in different medical fields, such as oncology^1^ and ophthalmology^2^. Recent advancements in light-responsive, biocompatible materials unlock new concepts, spanning from artificial vision^3^ to wireless stimulation of the nervous system,^4^ as well as tissue regeneration via phototherapy.^5^ In general, using exogenous functional materials to selectively manipulate cell activity is considered a less risky approach compared to optogenetics.^6^ Biology guided design principles have established intracellular and extracellular material interfaces able to photo-stimulate a range of biological systems. Successful examples include carbon-based nanomaterials^7^ and gold nanoparticles^8^ for photo-thermal cancer treatment, silicon nanowires for photo-modulation of neurons^9^ and other light-responsive silicon-based structures to control biological systems.^10^

Recently, organic photo-actuators — devices that use organic semiconductors able to modulate cell activity with light — attracted significant interest because of relatively simple fabrication requirements and device setup.^11^ For in vitro applications, organic semiconductor thin films are deposited on transparent conducting substrates and are directly interfaced with cells that are cultured on their surface. Organic semiconductors are highly biocompatible and show strong absorption of light in wavelengths spanning from ~400 nm to 700 nm. Therefore, light can be simply absorbed by the photosensitive organic semiconductor, and converted into a cue that modulates the activity of cells in close proximity. These unique advantages have been leveraged to build organic photo-actuators with high temporal and spatial resolution for both *in vitro*^12^ and *in vivo*^13, 14^ applications. Other than neurons, photo-actuators were also proved successful in controlling the behavior of non-electrogenic cells, such as astrocytes,^15^ endothelial colony-forming cells,^16^ and embryonic kidney cells (HEK-293) with light.^17^ Other applications of organic photo-actuators include the light-triggering the redox states of specific proteins,^18^ modulating the formation of reactive oxygen species in cell cultures,^19^ and controlling the intracellular redox equilibrium in model organisms.^20^

The exact mechanisms of controlling the activity of biological systems with organic photo-actuators can be quite diverse and are still not well understood.^21^ In general, organic semiconductors convert light into a cue that triggers cell reactions *via* three major mechanisms, i.e. photo-thermal,^12^ photo-capacitive^22^ and photo (electro) chemical^16^ stimulation. Photo-thermal stimulation usually occurs when photo-generated excitons in organic semiconductors — a strongly bound hole/electron pair^23^ — recombine, releasing their energy as thermal radiation. This mode of operation is usually happening when a single organic semiconductor is used as the photosensitive component and/or of prolonged exposure in high intensity light irradiation. The photosensitive component can be also made from a combination of p-type (hole transporting) and n-type (electron transporting) organic semiconductors, to form light sensitive p-n junctions.^24^ These p-n junctions are generally able to deliver large photovoltage and photocurrent at the polymer/cell interface due to exciton splitting that minimizes charge recombination. As a result, free electrons interact with cations in the electrolyte and induce a photo-capacitive modulation of the cell membrane.^25^ Free electrons can also react with oxygen or other electrolyte species to form reactive oxygen species and hydrogen peroxide. These photo-faradaic reactions can trigger ion channels on the cell membrane to modulate cell behavior. Overall, the photo-stimulation mechanisms involved in organic photo-actuators can be complex^26^ and specific for different biological systems. Eliciting light-sensitivity from a biologically relevant aqueous condition such as neuronal cells growing in culture would advance our understanding of biological systems without direct genetic manipulation (i.e., optogenetics) while further developing this technology.

Organic semiconductors can be molecularly tuned by chemical synthesis with unique absorption profiles and optoelectronic properties that can be leveraged to tune the sensitivity of photo-actuators at different light wavelengths^27^ and light intensities.^28^ The Głowacki group reported the development of organic p-n junctions based on conjugated small molecules and use them for the photo-capacitive stimulation of neuronal cultures^24^ and single cells.^29^ These devices are sensitive in the near infrared region of the optical spectrum and are able to photo-stimulate cells at low light intensities. Earlier studies from Lanzani and co-workers, used p-type poly(3-hexylthiophene) (P3HT) and the n-type fullerene derivative [6,6] phenyl-C61-butyric acid methyl ester (PCBM), to trigger action potentials in neurons upon light illumination.^30^ These results revealed the potential of organic semiconductors to be used in photo-stimulation of a wide range of biological systems. However, a number of both p-type and n-type organic semiconductors with improved optoelectronic properties have been recently developed for solid-state optoelectronic applications. In particular, non-fullerene electron acceptors (NFAs) have revolutionized the field of organic solar cells due to their excellent charge transport, strong optical absorption in the visible and NIR region and better energetic match with donor material.^31^ These materials show larger absorption coefficient, improved charge transport and lower exciton recombination rates compared with the materials used so far for photo-stimulation of biological systems. Therefore, the use of these newly developed materials and the systematic study of their operation in biologically relevant conditions will open new horizons for organic photo-actuators and potentially enable the photo-stimulation of biological systems at low light intensities.

Here we show the development of p-n junction photodiodes based on the p-type polymer poly[2,2′′′′ - bis [[(2-butyloctyl)oxy]carbonyl] [2,2′:5′,2′′:5′′,2′′′-quaterthiophene] −5,5′′′-diyl] (PDCBT) and the n-type polymer 3,9-bis(2-methylene-(3-(1,1-dicyanomethylene)-indanone))-5,5,11,11-tetrakis (4-hexylphenyl) – dithieno [2,3-d:2′,3′-d′]-s-indaceno[1,2-b:5,6-b′] dithiophene (ITIC). These photodiodes are made by simply spin coating the polymer solutions on a glass substrate support to form homogeneous PDCBT/ITIC bilayers with overall thickness of 55 nm. The p-n bilayer stack show good optical absorption in a wide range of wavelengths of the visible spectrum and good operational stability in aqueous electrolytes as verified by electrochemical quartz crystal microbalance studies. Controlled photo-electrochemical measurements, performed by spin coating the polymer films on conducting ITO substrates, reveal that relatively high photovoltage (>0.3 V) and high photocurrent (>0.6 mA/cm^2^) values can be produced when illuminated with low light intensity (~40 mW/cm^2^). An in depth photo-electrochemical analysis is presented and suggests photo-generated electron transfer to oxygen as the dominant process occurring at the polymer/electrolyte interface. The biocompatible nature of these materials is revealed by live/dead assays of different cell lines that are cultured directly on their surface (i.e. MDCKII, HEK293 and primary neurons). By simply spin coating the p-n bilayer junction on glass coverslips without the use of any conducting substrates or external electrodes we were able to control the activity of primary (mouse) cortical neurons with light. Several individual neurons were probed and frequent action potential were recorded when illuminated with 40 mW/cm^2^ white light. Identically cultured neurons on control glass or glass/photoresist layers show no action potential generation under the same illumination conditions, suggesting a non-photo thermal activation pathway. Under these conditions, photochemical oxygen reduction to H_2_O_2_ is the primary process. The evidence indicates that the action potential generation is driven by peroxide/reactive oxygen species (ROS) interaction with ion channels on the neuron membrane might be a possible general mechanism for neuron activation with light mediated by organic semiconductors.

## Results and Discussion

Constructing an efficient and stable p-n junction photodiode by using a layer-by-layer solution process is challenging. Since these devices are operating in an aqueous environment, two main film-processing conditions need to be carefully considered for optimum device stability: the solvent of choice to process the films and the adhesion of the p-n layers. To address these challenges, we first spin coated the p-type polymer PDCBT from a chlorobenzene solution, followed by spin coating the n-type non-fullerene acceptors (either ITIC and o-IDTBR) from toluene solution. This ensures that the solution processing do not damage the PDCBT film underneath. The chemical structures of the polymer used as well as the absorption profile of the PDCBT-ITIC polymer p-n junction is shown in figure 1. The cortical neurons used in this study were extracted from mouse embryos and cultured on the surface of these optically active platforms.

**Figure 1:**
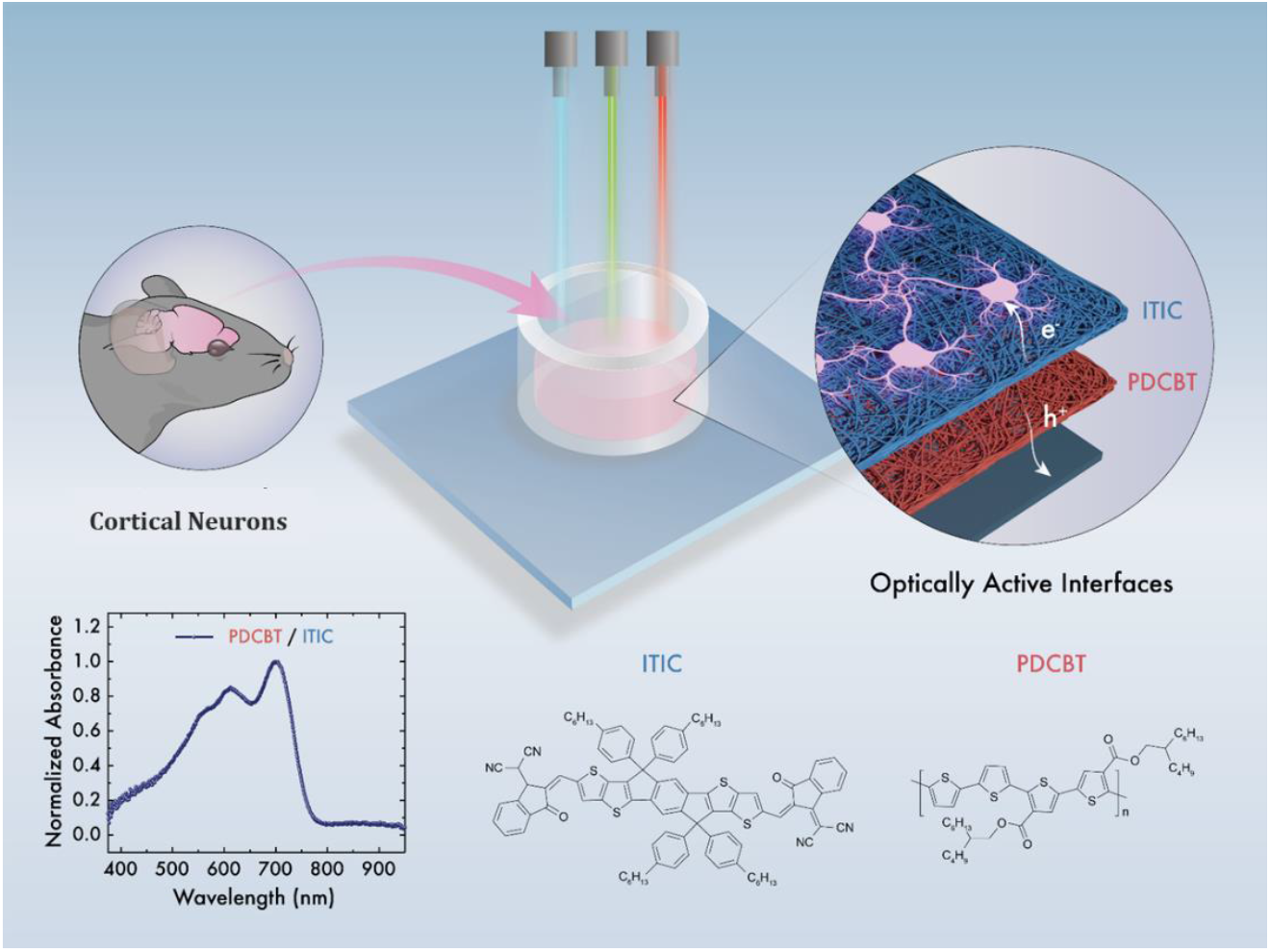
Schematic representation of the concept of photo-stimulation of neurons *In Vitro*. Mouse cortical neurons are extract and cultured directly on a photosensitive *P-N* polymer junction made by a simple layer-by-layer spin coating of the polymer. The absorption spectrum of the P-N junction as well as the chemical structures of the semi-conducting polymers used are shown in the bottom panel.

To understand the operation of our proposed p-n junctions in biologically relevant conditions we performed a series of photo-electrochemical measurements, which are summarized in Figure 2. First, we used electrochemical quartz crystal microbalance with dissipation monitoring (EQCM-D) equipped with a special module (figure 2a) where a window access allows for the simultaneous monitoring of mass changes and current upon illumination *in-operando*. We first coated the PDCBT polymer film directly on the QCM gold sensor and measured the thickness in both dry conditions and in PBS electrolyte, i.e. 27 nm and 28 nm respectively. We then spin coated the ITIC polymer film on the same sensor and its thickness was found to be 28 nm and 30 nm in air and in dry conditions respectively. The overall thickness of the p-n junction remains almost unchanged when immersed in PBS for several hours (i.e. 55 nm in dry conditions and 58 nm in PBS), indicating a stable “wet” operation of these p-n junctions – a major requirement to interface them with biological systems. After the p-n junctions were stabilized in PBS, a low intensity white light illumination was applied and the photocurrent response together with the *in-operando* mass change of the polymer films were recorder (figure 2c). Upon illumination, an average photocurrent of ~5 μA was recorded accompanied with almost no change in the mass of these polymer films during operation. This observation is indicative of a photo-electrochemical reaction at the polymer/electrolyte interface and opposes to a ion-uptake process which would manifest in a mass change.^32^

**Figure 2:**
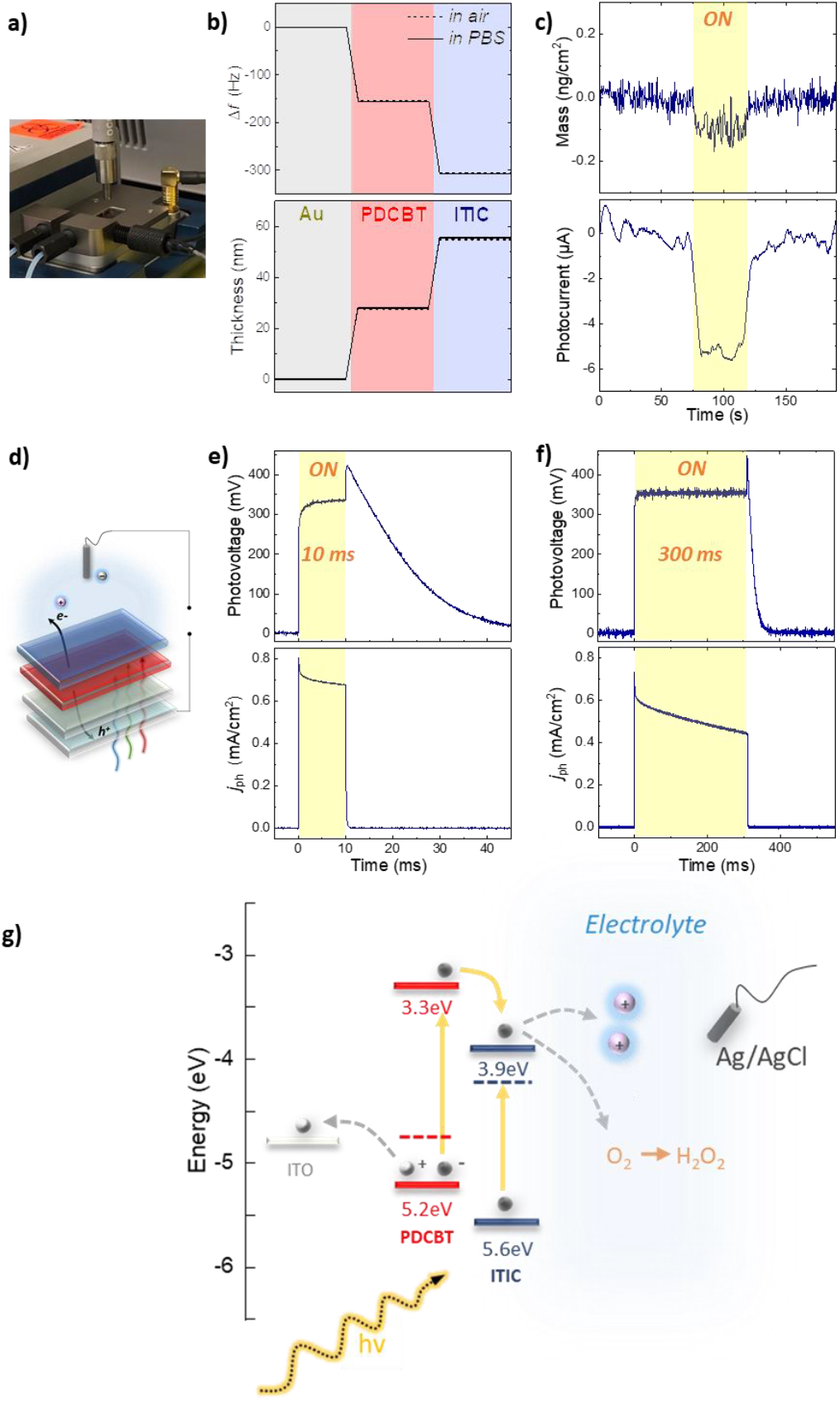
**a)** A digital photograph of the Photo-EQCM-D setup. The optical fiber on top is used to illuminated the donor-acceptor heterojunctions in the chamber through the window opening, while electrodes attached on the module record the photocurrent in addition to mass changes *in-operando***. b)** The thickness of the PDCBT-ITIC polymer films calculated by the raw QCM data (top graph) in dry conditions (solid line) and when immersed in the aqueous electrolyte PBS 1X (dashed line). **c)** Mass changes of the PDCBT-ITIC polymer films and the corresponding photo-current changes upon illumination *in operando.* **d)** Schematic of the setup used to study the photo-electrochemical operation of PDCBT-ITIC photodiodes in aqueous electrolytes. The photovoltage and photocurrent density response of the system is shown in **e)** for a 10 ms and in **f)** for a 300 ms light pulse of red light (λ_630nm_) with intensity of 33 mW/cm^2^. **g)** a relative energy diagram and the photo-electrochemical processes occurring after light exposure, showing photocapacitive charging at the semiconductor/electrolyte interface, as well as photocathodic charge transfer to oxygen, leading to hydrogen peroxide as the product.

Photo-electrochemical measurements with controlled light intensity and light pulse width were performed to establish the magnitude of photocapacitive and/or photofaradaic currents, which the p-n junctions can produce (Figure 2d–f). Both PDCBT and ITC where spin coated on ITO substrates and immersed in PBS with an Ag/AgCl as the counter electrode (figure 2d). The photo-current and photovoltage produced after illuminating the films with 10 ms and 300 ms light pulses are shown in figures 2e and 2f, respectively. In both cases we measured a photovoltage of >300mV and photocurrent density of >0.6 mA/cm^2^ when the photodiodes were illuminated at relatively low light intensity of 40 mW/cm^2^. These values are significantly larger that the values measured by single PDCBT or ITIC films (figure S1) indicative of the efficient exciton splitting at the PDCBT-ITIC interface and the formation of free electron carriers. The latter is verified by incorporating these p-n junctions in solid-state organic solar cell diodes (figure S2). The large open circuit voltage value obtained (i.e. 0.9 V) suggests efficient charge photo-generation and transport processes, arising from the optimum opto-electronic properties (e.g. charge mobility, optical absorption) of the materials used.^31^ It is therefore reasonable to expect that illuminated PDCBT-ITIC junctions can generate appreciable amounts of photocurrent and photovoltage in aqueous environments.

Further information for the mechanisms of the photo-charge generation by the polymer films operated in aqueous electrolytes can be extracted from photocurrent and photovoltage versus time graphs in figures 2e and 2f. Firstly, the polarity of the photocurrent is photocathodic, that is electrons are migrating to the semiconductor/electrolyte interface. We observe that both the voltage and the current are rapidly increasing and reach maximum values in the first few ms after illumination. This fast charging component can be ascribed to the photo-capacitive effect^25^ — free electrons are generated by the PDCBT-ITIC junctions and transported fast at the ITIC/electrolyte interface and interact electrostatically with positively charge ions in the electrolyte. This causes a capacitive displacement between the ITIC/electrolyte interface and the Ag/AgCl/electrolyte interfaces and is responsible for the fast voltage and current rise upon illumination. However, the high photocurrent and photovoltage values are sustained for the whole duration of the light illumination, indicative of a continuous faradaic reaction (pr reactions). Photocathodic current is maintained for over 90 minutes. Based on several previous works about the photocathodic behavior of organic semiconductor films, we assign this photocurrent to the oxygen reduction reaction.^33, 34, 35^ Those published studies indicate hydrogen peroxide as the dominant product of the photocathodic oxygen reduction (faradaic yield 90+%). To verify these observations we measured the H_2_O_2_ generation that is produced by the p-n junctions upon illumination (figure S3), and found a faradaic yield of 159% for photocurrent-to-peroxide conversion. Values above 100% are possible only if peroxide is produced via another pathway besides photocathodic charge transfer. An alternative pathway characterized in organic thin films is photochemical, where photogenerated excitons result in oxygen reduction with concurrent oxidation of some donor, often the organic photoactive material itself.^36^ To confirm that photochemical peroxide evolution indeed happens, we irradiated a film of the p-n junction on glass (no bias or current source), and found value 27 μM of H_2_O_2_ is produced, indicating a charge transfer process between photo-generated electrons and oxygen that leads to hydrogen peroxide generation via a photochemical mechanism.^33^ Photo-irradiation of PDCBT-ITIC junctions is therefore expected to lead to peroxide evolution.

The proposed mechanisms of operation are summarized in figure 2f. First, photons generated from the light source excite electrons located in the highest occupied molecular orbital (HOMO) of PDCBT and ITIC to the lowest unoccupied molecular orbital (LUMO). The HOMO and LUMO values were measured with ultra-violet photoelectron spectroscopy (UPS) (Figure S4) and the relative energy diagram is constructed as shown in figure 2f. We found that the LUMO of the n-type polymer ITIC provides an appropriate energetic step to accept the electrons that are photo-excited to the LUMO of PDCBT and as a result, the exciton splitting at this interface is efficient. These electrons are then transported to the ITIC/electrolyte interface and are transferred to molecular oxygen in the media to produce H_2_O_2_. When pristine PDCBT or ITIC films are used excitons recombine, to re-establish the low-energy ground state,^37^ leading to significantly smaller photocurrent and photovoltage values as well as smaller H_2_O_2_ generation upon illumination. Overall, the proposed PDCBT-ITIC junctions are highly photosensitive and can generate hydrogen peroxide both in photocathodic operation (with an underlying electrode and electrochemical circuit), or under photochemical operation, where neat films on insulating substrates photochemically reduce oxygen to peroxide.

We then used these photo-sensitive polymer interfaces as platforms to culture cells. We simply spin coated PDCBT and ITIC polymer films on glass coverslips and different types of cells were cultured on their surface. An important parameter of cell culture is the sterilization of the components that are used to grow cells to avoid cell contamination. Before cell seeding, the polymer films were sterilized by immersing them in 70 % ethanol solution for more than 1 hour. As shown in figure 3a, the ethanol sterilization process is not affecting the absorption spectra of both PDCBT and ITIC polymer films. This suggests that the optoelectronic properties of the polymer films are maintained. To evaluate the biocompatibility of PDCBT and ITIC polymer films we first studied the adhesion and growth of Madin-Darby Canine Kidney cells (MDCK II) on the polymer surface. As shown in figure S5, cells cultured directly on the sterilized polymer surfaces showed limited spreading on the surface and cell clusters were formed. This cell culture behavior is due to the hydrophobic nature of these polymer films as verified by the water contact angle measurements shown in figure 3b. To enhance cell spreading, we coated the polymer surfaces with thin layers of biological substances that are commonly used to facilitate cell adhesion to tissue culture such as collagen or Poly-D-Lysine (P-*d*-L). The formation of this adhesion layer reduce the water contact angle from 110 °C to 62 °C and from 109 °C to 58 °C for PDCBT and ITIC respectively (figure 3b) and as such, a healthy MDCK II tissue is grown on both polymer surfaces. The same approach was followed to grow healthy human embryonic kidney cells (HEK 293) on the surface of the PDCBT-ITIC films as proved by the live/dead assay measurements in Figure S6. These results prove the good biocompatibility of PDCBT and ITIC polymer films and show their potential to be used as platforms for different cell line cultures while maintaining their functional optoelectronic properties. Therefore, we used these platforms to grow cortical neurons extracted from mice. Figure 3c shows the live/dead assay images obtained from these neuronal cultures on the 14^th^ day after seeding. Almost all-individual neurons exhibit the formation of both soma and axons and a well-interconnected neuronal network is established, proving the excellent biocompatibility of these interfaces for neuronal cultures.

**Figure 3:**
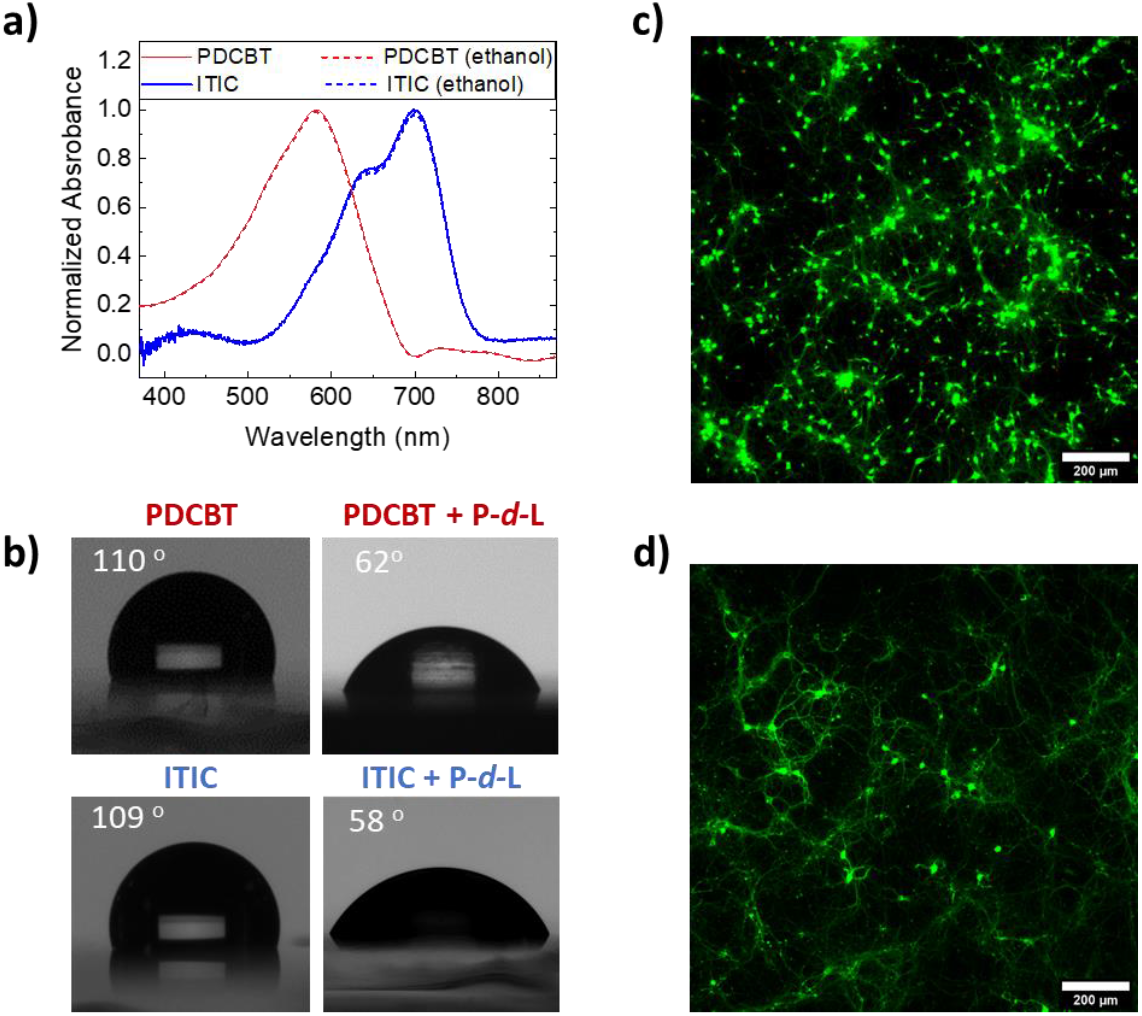
**a)** Optical absorption of the PDCBT (red) and ITIC (blue) conjugated polymer thin films as casted (solid lines) and after sterilization with 70% ethanol for 1 hour (dashed lines) **b)** Water contact angle of the PDCBT and ITIC polymer films before and after coating their surface with poly-d-lysine. Live/Dead assay of mouse cortical neurons on **c)** glass substrate and **d)** on glass substrate coated with PDCBT-ITIC thin films.

We then utilized the photo-sensitive properties of PDCBT-ITIC polymer films to test the possibility of photo-stimulating the cortical neurons cultured on their surface. It is important to note that the polymer films were directly spin coated on the glass cover slips without the presence of ITO or any other conducting substrate, therefore the experiment is designed to test for possible photochemical stimulation effects. Patch clamp measurements^38^ were performed and representative I/V traces of a mouse cortical neuron cell cultured on the PDCBT-ITIC coated coverslip are shown in Figure 4a. These traces confirm that the cells were capable of producing action potential. We then used a 10 second pulse of white light illumination at a low light intensity (40 mW/cm^2^) to excite the neuronal cultures, and at the same time, we monitored the resting membrane potentials (RPM) of individual neurons. As shown in Figure 4b, the resting membrane potential of an individual neuron that is grown on the PDCBT-ITIC surface shows a slow and progressive membrane depolarization which continues also after the light pulse ends, followed by a burst train of action potential spikes. This measurement is a clear indication that light stimulates neuron activity, most possibly due to the presence of the PDCBT-ITIC photo-sensitive layers that convert light into a cue. It is important to note that light illumination does not create any electrical stimulation artefact in the electrophysiological recording, allowing us to safely rule out any capacitive stimulation of the cell membrane. We performed a control experiment on individual neurons that are grown on top of pristine glass cover slips. These neurons do not respond to light stimulation even after a 20 seconds light pulse. We therefore eliminate the possibility of the direct interaction of light with the neurons without the mediation by the PDCBT-ITIC polymer films. We further investigated the mechanisms of photo-stimulation by growing neurons on a glass coverslip coated with a thin layer of photoresist. This photoresist is an insulating layer that absorbs light in similar wavelengths as the PDCBT-ITIC polymer junction and is effective in converting light into heat. No RPM change and action potential response was recorded for individual neurons grown on this film under the same illumination conditions (10 s of white light – 40 mW/cm^2^). This control experiment shows clearly that the generation of action potentials of neurons that are cultured on the PDCBT-ITIC polymer films is not likely due to any photo-thermal results since the same illumination conditions do not cause any neuron stimulation on the photo-resist. As shown in figure 4c, the depolarization of the RMP upon illumination occurs only on neurons that are grown on the PDCBT-ITIC junctions and not on the glass or photo-resist controls. A slight depolarization of the RPM is also observed on neurons that are grown on the individual p-type PDCBT layers or the n-type ITIC layers which is in line with the photo-electrochemical results observed in figure 2 and figure S1. The pristine PDCBT and ITIC films are less photo-sensitive compared with the p-n junctions and could explain why neurons grown on these pristine films are less affected by the light pulses compared with the neurons grown on the p-n junctions. Action potential generation has been observed by many individual neurons grown on several coverslips coated with PDCBT-ITIC polymer films as shown in figure 4d.

**Figure 4:**
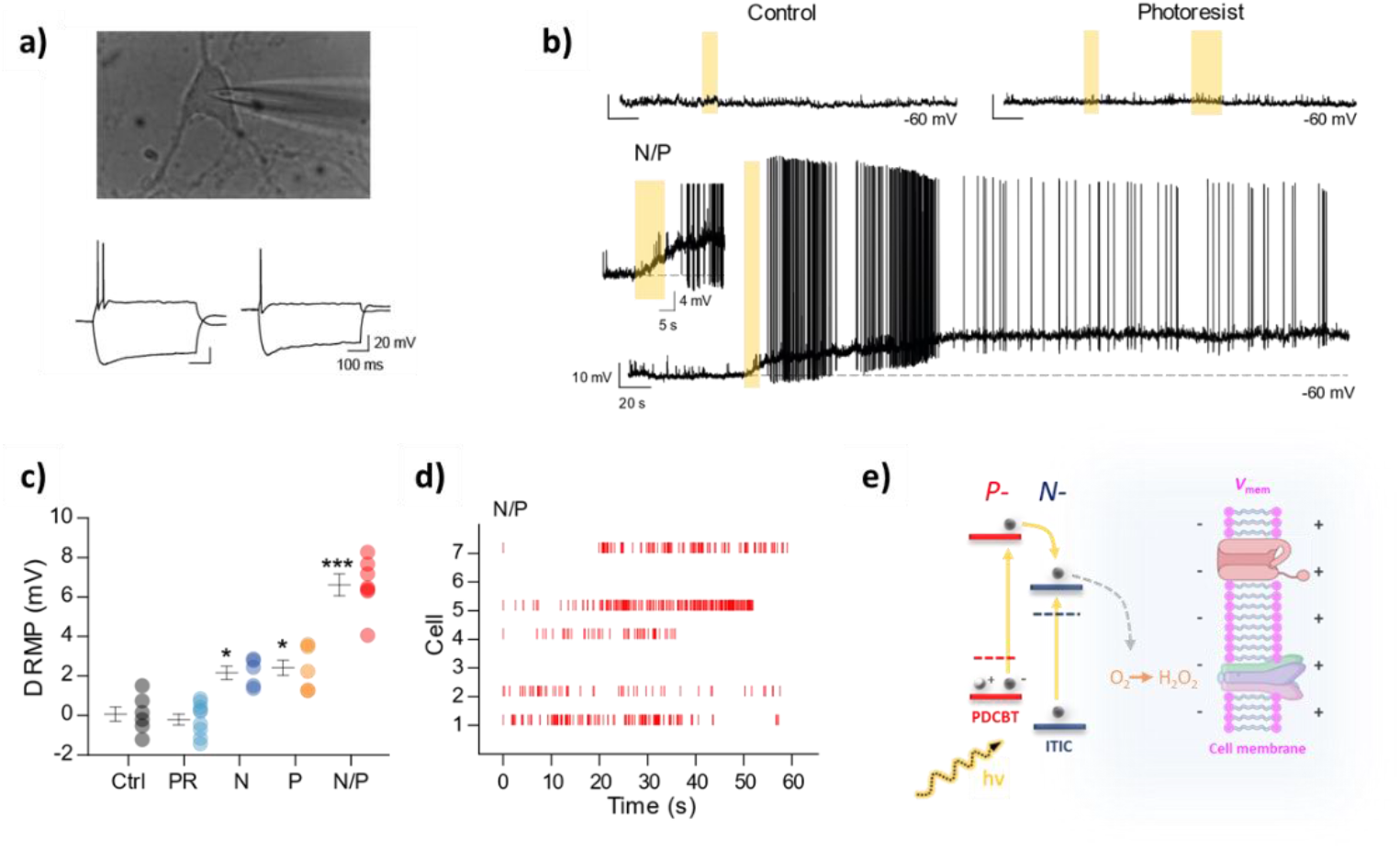
**a)** representative I/V traces of a mouse cortical neuron cell cultured on the PDCBT-ITIC coated coverslip showing that the cells are capable to produce action potentials. **b)** Representative traces in CC gap free mode in control and with the cells excited on the photoresist material (upper panels) and after the stimulation with light of a cell on the full device coated with the N/P polymers. **c)** Change in resting membrane potential of cells after 10 seconds of light exposure. **d)** Raster plots showing each spike evoked with the 10 seconds shinning of neurons seeded on N/P coated coverslips. **e)** Schematic of the proposed mechanism of photostimulation of neurons grown on top of the PDCBT-ITIC polymer p-n films.

We therefore suggest that PDCBT-ITIC junctions may serve as a biocompatible platform to mediate the stimulation of neurons with light. The evidence indicates that stimulation occurs through a photochemical pathway mediated by the polymer films. Since the devices are a neat organic film on an insulating substrate, we exclude any photocapacitive effect on the neurons. Since photothermal effects can be excluded, and it is clear that such films produce hydrogen peroxide upon illumination, we suggest that a photochemical stimulation mechanism with H_2_O_2_ is leading to the observed membrane depolarization and action potential stimulation. As shown in figure 4e, Transient Receptor Potential Vanilloid Receptor 1 (TRPV-1) channels may be responsible for the response. TRPV1 channels are endogenously expressed in many neuronal cell types, including rat cortical neurons.^39^ TRPV1 channels have recently been identified as suitable targets for photogenerated ROS stimulation. Transient Receptor Potential Vanilloid Receptor 1 (TRPV-1) channels^40^ in HEK 293 cells can be triggered by light stimulation mediated by organic semiconductors. TRPV1 channels are expressed in rat cortical neurons that form Ca^2+^ permeable cation channels and are activated by physiochemical stimuli, including pH change.^41^ H_2_O_2_ acts on this class of channels and have shown to produce excitatory effects on non-dopamine (DA) neurons in the midbrain where TRPV1 channels are also present.^42^ The same channels also seem to have a central role in the mechanisms of light stimulation of progenitor cells using organic semiconductors.^16^ Our electrophysiology recordings are consistent with a picture of TRPV1 mediated excitation, in particular the observation of a slow rise in RMP associated with light irradiation. Peroxide is produced by photo-irradiation, and this peroxide concentration steadily grows and activates nearby TRPV1 channels, leading to this slow membrane depolarization followed by bursts of action potentials. This excitation mechanism is schematized in Figure 4e.

## Conclusion

Solution processed heterojunctions made of the p-type polymer PDCBT and the n-type polymer ITIC (also known as non-fullerene acceptor) are proposed as highly sensitive photostimulation platforms of primary neurons *in vitro*. We found that these films convert light into cue through photo-electrochemical processes without the need of external electrodes or conducting substrates. Advanced analysis of the photocurrent and photovoltage waveforms combined with measurements of the amount of H_2_O_2_ that is produced, suggest efficient electron transfer between the polymers and molecular O_2_. These effects occur at the cell/polymer interface both mediated by the polymer photoexcitation and are responsible for triggering action potential on primary cortical neurons. As by-products of oxidative cellular metabolism, H_2_O_2_ and Reactive Oxidative species (ROS) are not only considered cellular toxic waste, but are key in regulating physiological conditions. New emerging evidence highlights their role in acting as signaling molecules controlling cell fate and guiding embryonic development, while possibly acting as neuromodulators. Although, the exact role of H_2_O_2_ and reactive oxygen species that might be produced are not fully understood, these stimulation pathways offer highly novel means to modulate cellular activity. However, their existence in excess may also involve toxic effects on cells and must be investigated in further details. Altogether, the simplified fabrication process together with the ability of wireless stimulation at low light intensity pave the way for the integration of these systems in multiplex bioelectronic devices. The biocompatibility of these materials with different types of tissue pave the way for the development of novel light therapies based on optically active organic semiconductors.

## Experimental

### Materials

glass-ITO substrates (sheet resistance 15 Ω/sq) were purchased from Xin Yan Technology LTD. PDCBT was purchased from 1-material inc. ITIC was purchased by ossila ltd. o-IDTBR was synthesized following the synthetic procedure reported by S. Holiday et al.^43^

### Device fabrication

Glass substrates fully coated with indium tin oxide (ITO) were cleaned by sonication in acetone and isopropanol, followed by oxygen plasma treatment. The p-type PDCBT layer was deposited from 10 mg ml^−1^ solution in chlorobenzene by spin coating at 2000 rpm resulting in a thickness of 40 nm. The solution was heated at 80°C prior to spin coating and spin coated hot to ensure good layer homogeneity. Both ITIC and o-IDTBR active layers were spin coated on top of PDCBT layers at 2000 rpm from 10 mg ml^−1^ solutions in toluene resulting in a thickness of 40 nm. Organic solar cells devices were fabricated with a p-n device architecture (NiO/PDCBT/ITIC or o-IDTBR/Al). NiO was prepared using a sol-gel method on ITO substrates reported elsewhere followed by the sequential deposition of the p-n layers. Finally, 100nm thick aluminum layer was thermally evaporated on top of the active layers of the devices through a shadow mask yielding active areas of 0.045 cm^2^ in each device.

### Characterization

Measurements of photovoltage and photocurrent of the p-n junction based devices were conducted inside a dark Faraday cage and the electrophotoresponse (EPR) data were collected via a high resolution 15-bit two-channel PicoScope 5243B oscilloscope as described previously.^29^ Briefly, the backside ITO of the OEPC was contacted with a probe electrode connected to the positive terminal of an oscilloscope. Meanwhile, the negative terminal was connected to an Ag/AgCl electrode in 0.1 M KCl electrolyte, making contact to the top of the organic layer. The intensity of the LED used (wavelength = 630nm) is 0.33 mW mm^−2^. Local H_2_O_2_ evolution was measured in situ using a 4-channel micro-amperometric amplifier system (TBR4100, World Scientific Instruments), with 4-channel ADC board (LabTrax, World Scientific Instruments). The respective sensor probe used was ISO-HPO-2. The sensor was kept constantly polarized at a bias of 450 mV and calibrated before the measurement following the procedure reported in the instruction manual. The increase in H_2_O_2_ concentration was recorded using a LabScribe software (World Scientific Instruments). The experiment has been carried out placing the sample in a PBS containing vial and shining white light based LED of 942 mW optical power for 3h afterward the microamperometric sensor (Clark electrode) has been placed in the vial to detect the amount of H_2_O_2_ evolved after 3h of reaction. The concentration of H_2_O_2_ evolved after the reaction was of 27 μM. Longer measurements of photocathodic behavior shown in Figure S3 were conducted using an Ivium Vertex One potentiostat and a three-electrode sealed photo-electrochemical cell, with peroxide concentration quantified using a photometric assay with tetramethylbenzidine as described in Ref.^44^ To measure thickness of the conjugated polymer thin films a Dektak-Veeco stylus profilometer was used. Current density versus voltage (*J–V*) characteristics of the solar cells under study were measured by using a Xenon lamp at AM1.5 solar illumination (Oriel Instruments) calibrated to a silicon reference cell with a Keithley 2400 source meter and a custom made LabVIEW software. Optical absorption profiles were recorded with an Ocean Optics QE65 Pro Spectrometer, Ocean Optics HL-2000-FHSA halogen light source, OceanView software, and a sample holder purchased from redox.me (MM Spectro-EFC). Ultraviolet photoelectron spectroscopy (UPS) measurements were recorded using a SPECS PHOIBOS 100 hemispherical electron energy analyzer in a custom built ultrahigh vacuum (UHV) system with a base pressure of ~1 × 10^−10^ mbar. Samples were excited at 21.21 eV using a He I plasma source. UPS measurements were perform on polymer films spin coated on a gold foil identically with the ones used for the photo-stimulation experiments. Water contact angle measurements were performed with a KRUSS instrument using the sessile drop method.

### Electrochemical quartz crystal microbalance with dissipation monitoring

We performed EQCM-D measurements using a Q-sense analyzer (QE401, Biolin Scientific). Thickness and swelling measurements were performed as follows. First, we recorded the QCM-D response of the bare Au sensors in the air, followed by the injection of the PBS 1X solutions into the chamber. This resulted in large shifts in frequency (*f*) and dissipation of energy (*D*), due to the density differences between the two media. The measurements were then stopped, the sensors were removed and the PCDBT polymer film was spun cast directly on the same sensor from a 10 mg/mL hot chlorobenzene solution at 2000 rpm. The absolute *f* value for the polymer coated sensor was obtained both in air and in PBS 1x, after the *f* signal was perfectly flat (i.e., f <0.5 Hz) assuring that the system is in equilibrium. The sensor was then removed again, ITIC film was spun cast from a 10 mg/mL toluene solution at 2000 rpm, and inserted back in the chamber to measure the absolute *f* in air and in PBS 1X. We then compared the absolute difference in *f* for multiple overtones between the bare sensor and the polymer coated sensor, both in air and in PBS 1X by using the function “stitched data” of Q-soft software. This function compares the selected datasets based on the raw frequencies measured and excludes the effect of the different densities between the two media. Thus, the difference of the *f* values of the stitched data is directly analogous to the thickness of the polymer in both media, which is calculated by using the Sauerbrey equation:

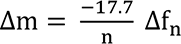

Q-Tools and D-find software were used for the modelling and data analysis. After the swelling measurements, an Autolab PGstat128N potentiostat was coupled with Q-sense electrochemistry module and the current was recorded. The three-electrode setup was comprised of Ag/AgCl reference, Pt counter and Au/polymer EQCM-D sensor as the working electrode. An optical fiber was attached on the window of the module and white light pulses were applied with an Ocean Optics HL-2000-FHSA halogen light source. The current was recorded with the potentiostat simultaneously with the raw frequency changes that were recorded with the QCM and therefore, the changes in the thickness (or mass) of the polymer films as well as the photocurrent were able to be extracted *in-operando*.

### Cell Culture

Human Embryonic Kidney cells (HEK 293) and Madin-Darby Canine Kidney cells (MDCK II) were cultured in DMEM supplemented with 10% fetal bovine serum, 2 mM Glutamax and 0.5% PenStrep 100X (10 000 U.mL^−1^ Penicillin, 10 000 μg.mL^−1^ Streptomycin). All reagents were purchased from Invitrogen. Cells were routinely maintained in incubators at 37°C and 5% CO_2_. Devices were sterilized using 70% ethanol for 30min and rinsed with water prior seeding. Primary cortical neurons were collected from mice. They were handled in accordance with the animal research advisory committee guidelines of the National Institutes of Health (NIH), and procedures were approved by the Institutional Animal Care and Use Committee (IACUC) of King Abdullah University of Science and Technology (KAUST). Primary culture of cortical neurons was prepared from embryonic day 17 CD-1 IGS mouse embryos (Charles River, UK). Cerebral cortical tissue was dissected, collected and incubated 30 min at 37°C in Hank’s Balanced Salt Solution (HBSS) supplemented with papain (20 units.mL^−1^; Worthington Biochemical), L-127 cysteine (1 mM; Sigma) and DNase I (5 mg.mL^−1^; Worthington Biochemical). Cortical cells were mechanically dissociated and filtered using a 40 μm cell strainer (Fischerbrand) and centrifuged for 5 min at 200xg. The supernatant was carefully removed, and cells were re-suspended in Neurobasal medium (Gibco) completed with B27 supplement (Gibco), 500 μM L-glutamine (Sigma), and 0.5x penicillin/streptomycin solution (Gibco). 20 μL of media containing 10^5^ cells were seeded on top of the device and incubated between 15 and 30 min to the neurons settle. Then, 1 mL of warm (37°C) cell media was carefully added to the well. After 4 h, the media was replaced to remove dead cells. Neurons were incubated at 37°C, 95% humidity, and 5% CO_2_. Media changed twice a week and were replaced with half volume of a fresh batch.

### Live/Dead assay

Cells were seeded on top of the device and incubated for 24-48 h for epithelial cells and 14 days for cortical neurons. Then media is removed and a mixture of calcein-AM, propidium iodide (Sigma-Aldrich) is applied to cells. After 5-15 min of incubation, devices were rinsed three times with HBSS (Invitrogen), and cells were imaged using an inverted fluorescent microscope (DMi8, Leica Microsystems).

### Calcium Imaging

Fluo-4,AM (F14201, Invitrogen) was used to observe the calcium fluctuation inside the cells. To prepare the Fluo-4,AM stock solution, 50 μg of Fluo-4,AM were dissolved with 22.8 μL of DMSO. Then 2 μL of Fluo-4,AM stock solution was mixed with 2 μL of 20% Pluronic F-127 (P3000MP, Invitrogen) and 996 μL of HBSS solution, giving a final concentration of Fluo-4,AM of 2 μM. 500 μL of this solution was then added to the cell culture which was incubated at 37°C for 30 min. The neurons were gently washed 3 times with HBSS solution and incubated at 37°C for another 30 min before we performed imaging measurements with an inverted fluorescent microscope (DMI8, Leica Microsystem).

### Patch Clamp

Primary cortical neurons were visualized using an inverted microscope Carl Zeiss Axio Observer.A1 (Carl Zeiss, Oberkochen, Germany). Neurons with classical pyramidal shape were chosen to perform the experiments, it is convenient to clarify that the experiments were done under minimal light conditions in a dark room to prevent the activation of the photosensitive devices before patch-clamping the cells. Patch pipettes were pulled from borosilicate capillaries using a P-1000 Flaming Brown puller (Sutter Instruments) to a final resistance of 4-5 MΩ when filled with a solution with the followed composition (in mM): K^+^-Gluconate 135, KCl 10, EGTA 1, CaCl_2_ 0.1, HEPES 10, Mg-ATP 5, Na_2_-GTP 0.5; pH=7.2-7.3 adjusted with HCl. Current clamp recordings were done at room temperature in normal HEPES-based solution with the followed composition (in mM): NaCl 135, KCl 2.5, CaCl_2_ 2, MgCl_2_ 1, HEPES 10, glucose 10; and were obtained with an Axopatch 700B amplifier (Axon Instruments, Molecular Devices, Sunnyvale, CA) and digitized at 20 kHz and low-pass filtered at 10 kHz (Digidata 1550, Molecular Devices). All data were acquired and analyzed offline with the pClamp 10.7 software (Molecular Devices). Once the gigaseal is broken and the whole cell configuration is achieved, we recorded the resting membrane potential (RMP) and ran an I/V curve by injecting square current pulses (from −200 in 50 pA steps, 1 sec) until elicit at least one action potential. Cells with RMP more depolarized than −50 mV were discarded from this study. A gap-free protocol was continuously recorded, and after a stable baseline, the light (intensity 40 mW/cm^2^) was turned-on for 1, 2, 5, 10 and 20 seconds to directly illuminate the patched neuron.

## Author Contributions

A.S. conceived the research, fabricated the devices, performed EQCM-D, optical absorption, photo-electrochemical, contact angle measurements, assisted on biocompatibility studies and wrote the manuscript. A.H. performed cell culture, calcium image and live/dead assays with assistance from N.S. and M.K. G.H.L, H.F. and P.M. performed patch clamp measurements and analyzed the results. N.G., D.B. and I.M. assisted on device fabrication and selection of materials. L.M. and E.D.G. performed photoelectrochemical measurements and peroxide quantification. A.S., E.D.G. and S.I. designed the experiments and supervised the work. All authors discussed the results and assisted in manuscript input.

## Acknowledgements

A.S. acknowledges funding from the European Union’s Horizon 2020 research and innovation program under the Marie Skłodowska-Curie grant, MultiStem (No. 895801). We thank Marios Neophytou, Akmaral Seitkhan, Joel Troughton for fruitful discussions and assistance in device fabrication. We also thank Professor Thomas Anthopoulos and Professor Róisín M. Owens for fruitful discussions. Antonio García, scientific illustrator at KAUST, created figure 1.

## Supporting Information

**Figure S1:**
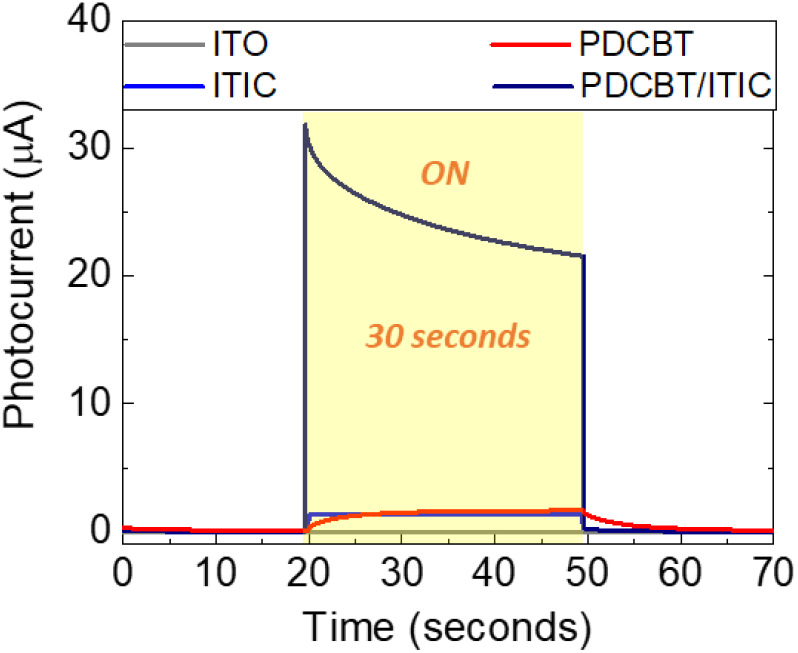
Photocurrent measurements of ITO only (grey line) ITO/PDCBT (red line) ITO/ITIC (blue line) ITO/PDCBT/ITIC (dark blue line) upon illumination with white light at 40 mW/cm^2^.

**Figure S2:**
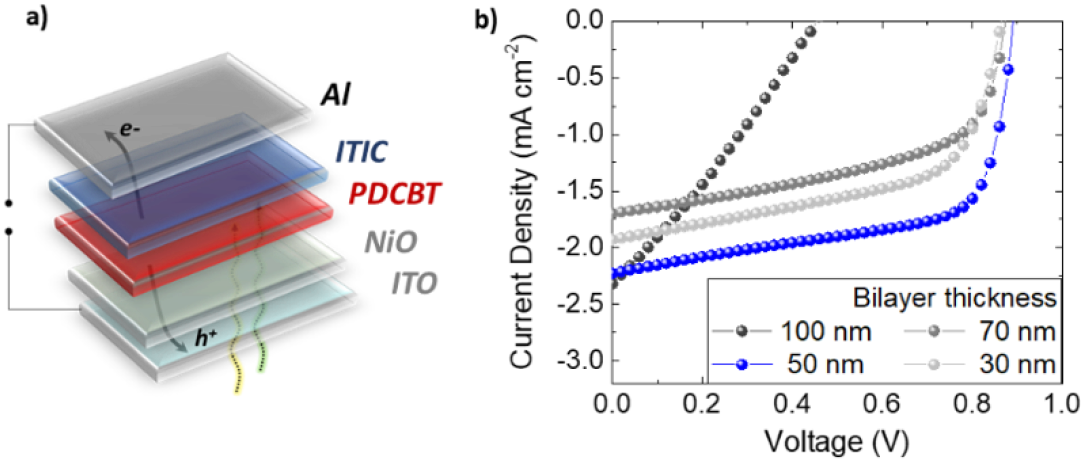
**a)** Schematic of the bilayer organic solar cell devices used - ITO/NiO/PDBCT/ITIC/Al. **b)** Current density versus voltage characteristics of the organic solar cells measured with different overall PDCBT-ITC thickness. The maximum short circuit current (2.2. mA/cm^2^) as well as the maximum open-circuit voltage (0.9 V) is achieved with a total PDCBT-ITIC thickness of 50 nm.

**Figure S3:**
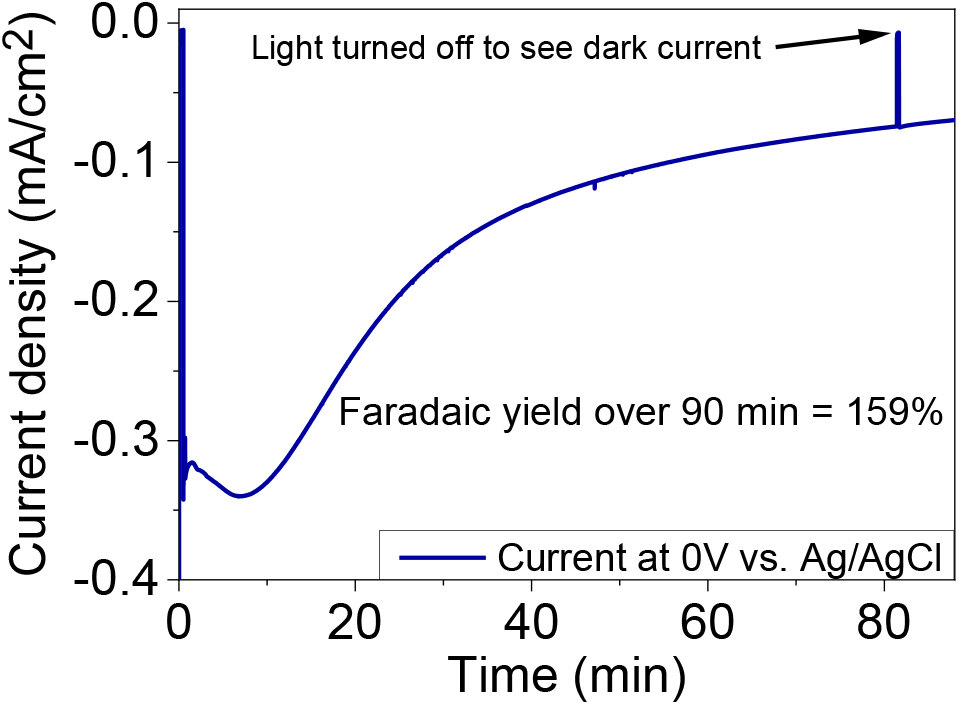
Photo(electro)catalysis of a representative sample ITO/NiO/PDCBT/ITIC illuminated with a halogen lamp as light source; E[V] = 0 Vt = 5400s.

**Figure S4:**
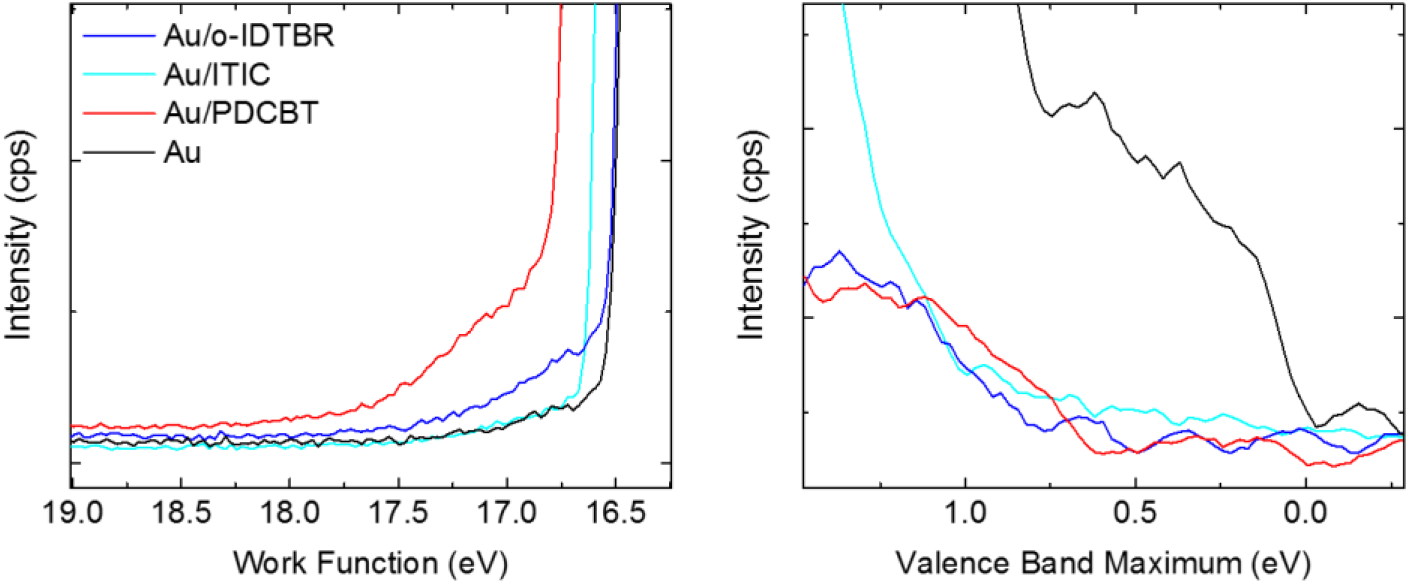
Ultraviolet photoelectron spectroscopy measurements of the polymer semiconductors thin films used in this study, i.e. PDCBT (p-type), ITIC (n-type) and o-IDTBR (n-type).

**Figure S5:**
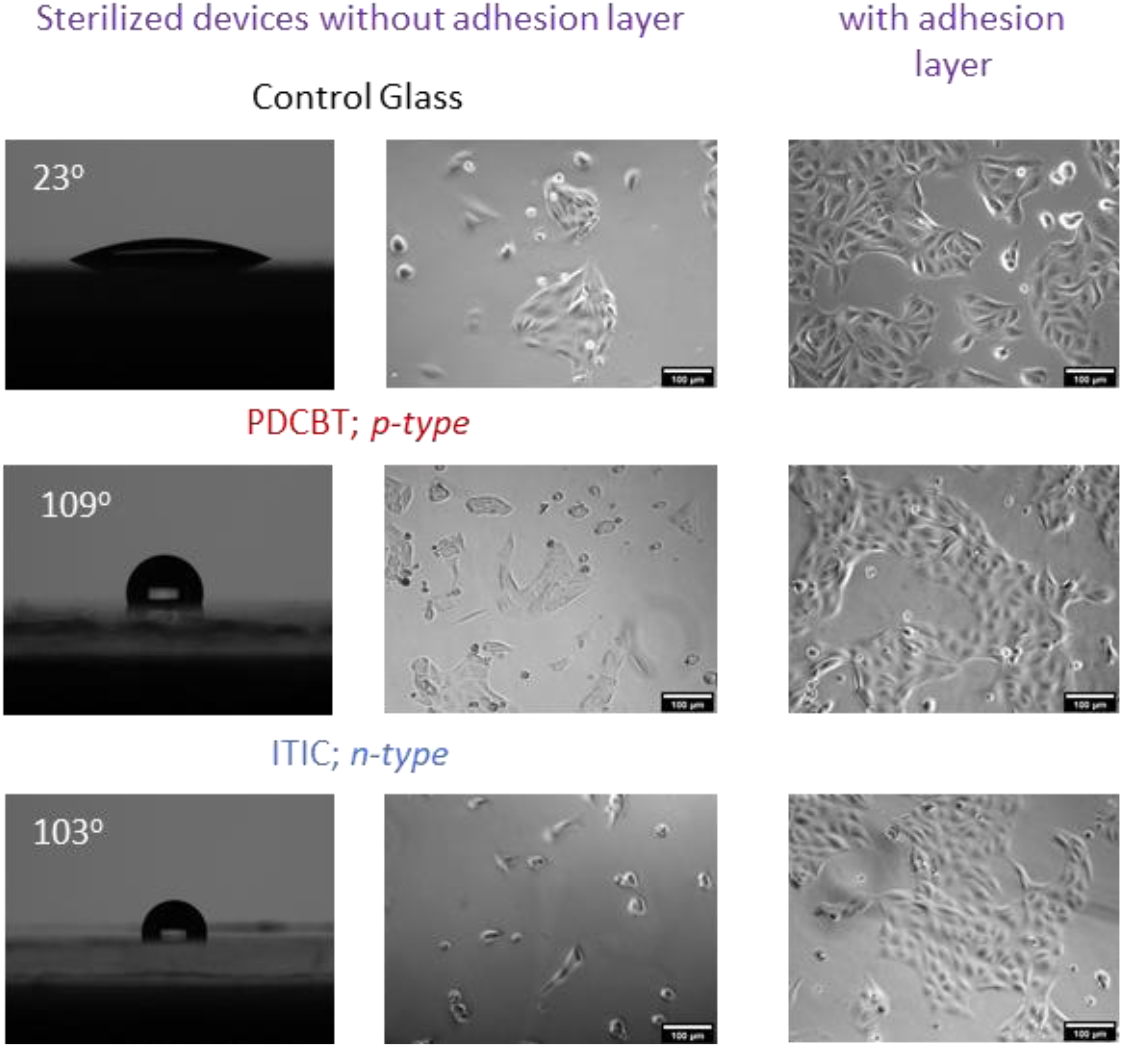
Madin-Darby Canine Kidney cells (MDCK II) cells cultured on the surface of control glass substrates (top raw), on PDCBT surface (middle raw) and on ITIC surface (bottom raw). The use of a thin adhesion layer (rat tail collagen) improves cell adhesion on the polymer surfaces and the formation of an epithelial layer. Scale bars = 100 μm.

**Figure S6:**
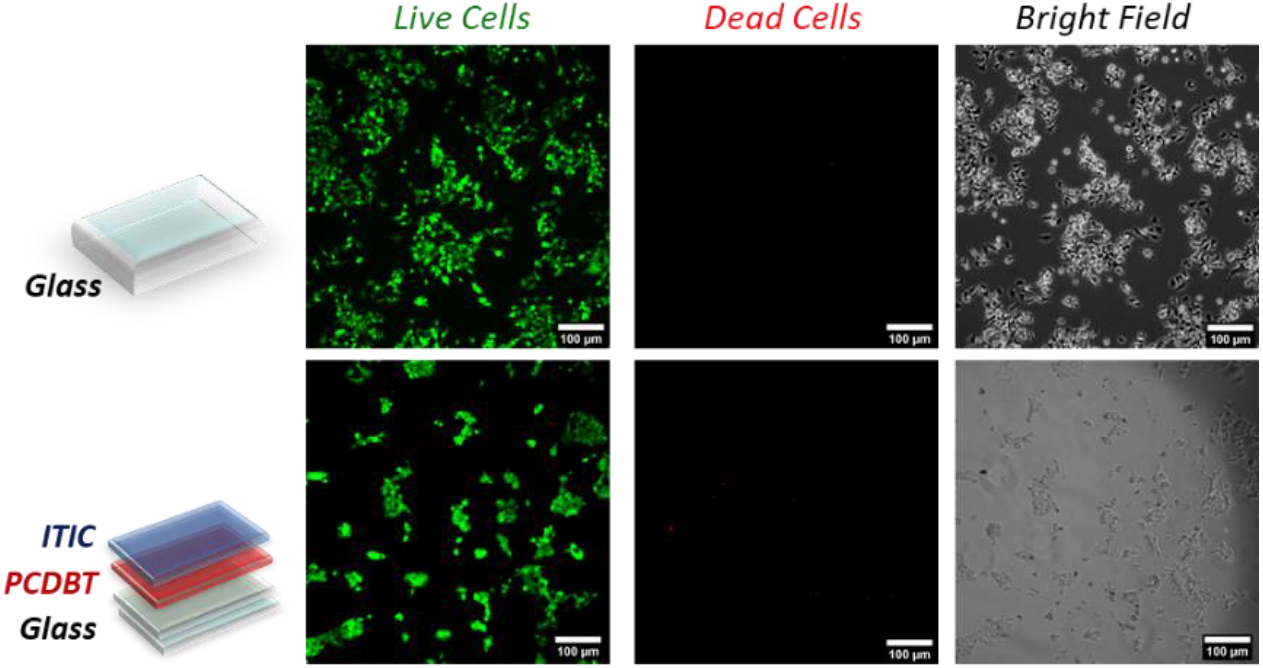
Live/dead assay of Human Embryonic Kidney cells (HEK 293) cells cultured on the surface of control glass substrates (top raw), and on PDCBT/ITIC surface (bottom raw). Scale bars = 100 μm.

